# Experimental warming influences species abundances in a *Drosophila* host community through direct effects on species performance rather than altered competition and parasitism

**DOI:** 10.1101/2020.12.22.423937

**Authors:** Mélanie Thierry, Nicholas A. Pardikes, Chia-Hua Lue, Owen T. Lewis, Jan Hrček

**Author notes:** **Author contributions**, MT, OTL and JH conceived the project; MT, NP and C-HL collected the data; MT analyzed the data. All authors contributed critically to the drafts and gave final approval for publication.

## Abstract

Current global warming trends are expected to have direct effects on species through their sensitivity to temperature, as well as on their biotic interactions, with cascading indirect effects on species, communities, and entire ecosystems. To predict the community-level consequences of global change we need to understand the relative roles of both the direct and indirect effects of warming. We used a laboratory experiment to investigate how warming affects a tropical community of three species of *Drosophila* hosts interacting with two species of parasitoids over a single generation. Our experimental design allowed us to distinguish between the direct effects of temperature on host species performance, and indirect effects through altered biotic interactions (competition among hosts and parasitism by parasitoid wasps). Although experimental warming significantly decreased parasitism for all host-parasitoid pairs, the effects of parasitism and competition on host communities did not vary across temperatures. Instead, effects on host relative abundances were species-specific, with one host species dominating the community at warmer temperatures, independently of parasitism and competition treatments. Our results show that temperature shaped a *Drosophila* host community directly through differences in species’ thermal performance, and not via its influences on biotic interactions.

## Introduction

It is becoming evident that many species are declining as the climate changes [1], and increasing numbers of extinctions are expected as a result in the coming decades [2]. Animals are directly impacted by warming temperatures through changes in their fecundity, mortality, metabolic rates, body growth rate, and phenology [3–6]. Species in the tropics are likely to be more sensitive to global warming because they are closer to their upper thermal limits [2,7], and the predicted increase in temperatures by a few degrees would exceed their thermal maxima. Ectotherms, such as insects, have particularly narrow thermal limits and are facing severe declines in abundances with rising temperature [8]. Warming temperatures directly affect physiology and demography depending on species’ thermal tolerances (i.e., their ability to survive exposure to extreme temperatures) and their thermal performance (i.e., their fitness-related traits over a range of temperatures). Both thermal tolerance and thermal performance are expected to influence population sizes and community structure with ongoing global warming [4].

However, ecological communities are not defined solely by the species that compose them, but also by the way those species interact with one another, via both trophic and non-trophic interactions [9,10]. Trophic interactions, such as predation, herbivory, or parasitism have strong effects on community composition and evenness [11,12]. Non-trophic interactions such as competition and pollination are also ubiquitous and can alter community composition in many ways (e.g. if some species are competitively excluded, or if species coexistence is enhanced) [13–15]. Trophic and non-trophic interactions act together to structure ecological communities [16–18], and a theoretical understanding is emerging of how these different types of interactions shape the structure and dynamics of more complex ecological networks [19]. However, empirical evidence from terrestrial and species-rich communities remains sparse, especially in a climate warming context. Warming temperatures are expected to have direct effects on both component species and their interactions [20,21]. Temperature can alter resource-consumer interactions via its effects on metabolic processes such as growth and reproduction, and change in behaviors [22–24]. The main mechanisms behind species interaction response to climate change are the differences in effects among interacting species, such as asymmetrical responses in their phenology [25], growth rate [26], and body mass [27]. Furthermore, species interaction changes with warming temperatures can have indirect cascading effects on individual fitness, populations and communities [24,28,29]. Despite calls for more investigations of how species interactions respond to global climate change [30,31], most such studies focus either on aquatic systems [20,32], on a single interaction type [33], or on a small number of species [34]. We urgently need more data to predict how environmental changes modify different types of interactions (both trophic and non-trophic) in more complex ecological networks [35,36].

Insect host-parasitoid communities are excellent model systems to investigate how species and their interactions respond to warming temperatures [13]. Parasitoids are insects which develop in or on the bodies of arthropod hosts, killing the host as they develop, and playing an important role in regulating host populations in both natural and agricultural ecosystems [37]. As ectotherms, many parasitoid traits involved in species interactions are sensitive to changes in temperature [38,39]. Empirical studies suggest that global warming could weaken top-down control by parasitoids by increasing parasitoid mortality, by decreasing parasitoid virulence and/or increasing host immune response, and by increasing host-parasitoid asynchrony, thus increasing the frequency of pest outbreaks [40–42]. However, most studies of host-parasitoid interactions are limited to a pair of interacting species, and it is unclear how host-parasitoid communities respond to warming temperatures when more complex systems are considered [13,43]. Parasitoids can mediate host coexistence, but the outcome may depend on temperature [44]. Moreover, competitive interactions among hosts can affect the responses of species and communities to environmental changes [29], but such responses may differ for intraspecific and interspecific competition [45]. Thus, to help forecast the impacts of global warming on host-parasitoid communities, it will be critical to examine the combined responses of species and their interactions under simulated warming conditions.

In this study, we use a laboratory experiment to investigate how temperature affects host communities (host relative abundances) directly through difference in species responses, and indirectly through effects on parasitism and host competition. We measured host body mass to provide a further proxy for host fitness under the different treatments. We focus on a set of three *Drosophila* species and two parasitic wasp species which form part of a natural *Drosophila-parasitoid* community in Australian tropical rainforests. We test the predictions that elevated temperature will have both direct effects on host relative abundances depending on the species thermal performances, and effects on their interactions by changing both hosts competitive abilities and parasitoids virulence [38], with indirect cascading effects on host relative abundances in the community [13]. Effects of temperature on host competitive interactions will be linked to species thermal performance, while effects on parasitism will be linked to effects of temperature on parasitoid attack rate and virulence. An interactive effects of trophic and non-trophic interactions on host relative abundances is expected due to a trade-off between resistance to parasitoids and larval competitive abilities [46]. This study provides an important step in disentangling the direct and indirect effects of warming on structuring tropical *Drosophila* communities, which will be critical for predicting effects of climate change on tropical insect communities.

## Materials and Methods

### Study system

The experiment was established from cultures of *Drosophila* species and their associated parasitoids collected from two tropical rainforest locations in North Queensland, Australia: Paluma (S18° 59.031’ E146° 14.096’) and Kirrama Ranges (S18° 12.134’ E145° 53.102’) (<100 m above sea level). *Drosophila* and parasitoid cultures were established in 2017 and 2018, identified using both morphology and DNA barcoding, and shipped under permit to the Czech Republic. Three host species (*Drosophila birchii, D. pseudoananassae* and *D. sulfurigaster,* together accounting for ~ 48% of the host communities at the study sites [47]) and two of their natural larval parasitoid species *Asobara* sp.1 (Hymenoptera: Braconidae; Smithsonian National Museum of Natural History (NMNH) reference vouchers USNMENT01557096 and USNMENT01557091) and *Leptopilina* sp.1 (Hymenoptera: Figitidae; NMNH reference vouchers USNMENT01557104 and USNMENT01557109) able to parasitize all three host species were used in this experiment [note: these are new undescribed species of parasitoids and we will link this manuscript with a taxonomic paper during revision and include GenBank and BOLD accession codes for reference sequences for these species to clearly define their identity]. Data on thermal performance of the three host species are available [48,49]. All cultures were maintained at 23°C on a 12:12 hour light and dark cycle at Biology Centre, Czech Academy of Sciences. *Drosophila* isofemale lines were maintained on standard *Drosophila* medium (corn flour, yeast, sugar, agar and methyl-4-hydroxybenzoate) for approximately 15 to 30 non-overlapping generations. To revive genetic variation, five lines from each host species were combined to establish mass-bred lines immediately before the start of the experiment. Isofemale lines of parasitoid lines were maintained for approximately 10 to 20 non-overlapping generations prior to the start of the experiment by providing them every week with 2-day-old larvae of *Drosophila melanogaster*. This host species is not present naturally at the field locations where hosts and parasitoids originated, and was not used in the experiment, thus avoiding bias of host preferences. Single parasitoid isofemale lines were used.

### Experimental design

To disentangle the effects of warming temperatures on host species and their interactions, we manipulated the presence of parasitoids and interspecific competition between host species in a fully factorial design (Figure 1) at ambient and elevated temperatures. As the focus of the experiment was to compare the direct and indirect effects of warming temperatures on host communities, competitive interactions between parasitoids were not assessed nor manipulated, but potentially present in all treatments with parasitoids. Parasitoid preferences were not quantified, but the two parasitoid species used were able to parasitize all three hosts species during trials. Transparent plastic boxes (47cm x 30cm x 27.5cm) with three ventilation holes (15 cm in diameter) covered with insect-proof nylon mesh served as the experimental units. Each box contained three 90 mm high and 28 mm diameter glass vials containing 2.5 mL of *Drosophila* food medium. Interactions were manipulated by establishing single-species (Figures 1a and 1c) or mixed-species (Figures 1b and 1d) vials, and by including (Figures 1c and 1d) or excluding (Figures 1a and 1b) parasitoids. A total of 60 three-day-old virgin adult hosts, with 1:1 sex ratio, were placed in each vial for 48 hours to allow mating and oviposition (i.e., a total of 180 adults per box). In the multi-host treatment, the 60 hosts were split evenly across the three species (i.e., 20 adults for each species). The density of adult hosts was selected based on preliminary observations to achieve a high level of resource competition (i.e., the density at which strong intraspecific competition was observed for all host species; Supplementary Table S1) while keeping the number of adults for each of the three host species and the total number of adult hosts consistent across treatments and species. The treatment allowed competition both at the adult stage for oviposition space, and at the larval stage of their offspring for food resources [50,51], but we did not aim to identify which was the primary source of competition. All results relate to the host offspring (their abundances and frequencies). For treatments that included parasitoids (Figures 1c and 1d), ten parasitoids (3-7 days old, 1:1 sex ratio) from each species (n = 2, i.e. 20 parasitoids per box), corresponding to 9 % of the total number of adult hosts, were placed in a box for 72 hours, creating high but realistic parasitoid pressure (within the range of parasitism rate observed in this system in nature: 8-42% [47]). We aimed to study the impact and interaction of parasitism and host competition when both are strong, but realistic, to detect any effect. Vials were removed from the boxes simultaneously with the parasitoids and individually sealed. Each treatment was replicated once across four time-blocks, and each treatment and replicate were therefore represented by three vials. The duration of the experiment corresponded to a single generation of both the hosts and the parasitoids. Each set of treatments was replicated once during a single day, and was repeated over four days (i.e., blocks).

**Figure 1.**
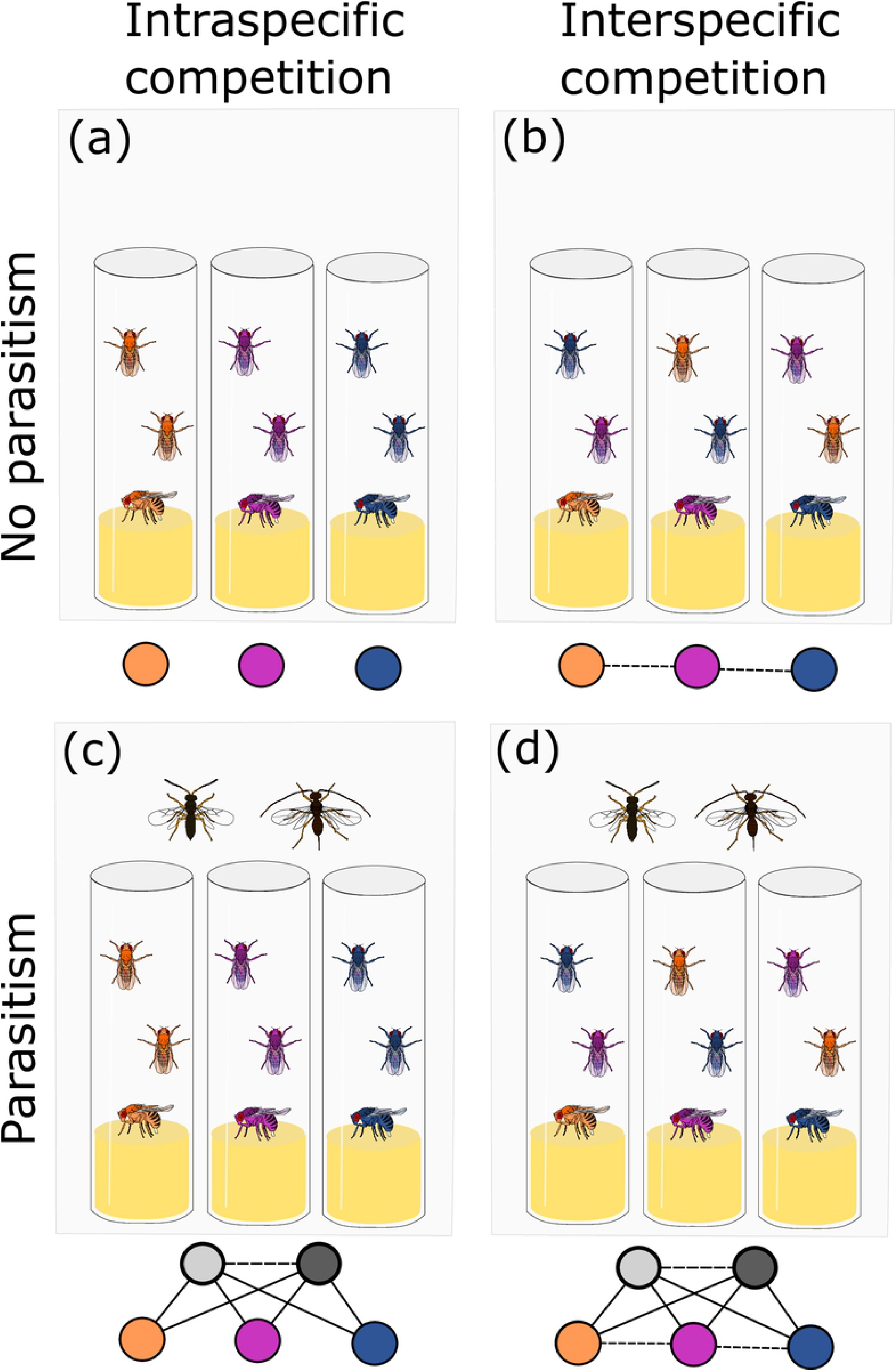
Schematic representation of experimental treatments. Orange, pink, and blue nodes represent the three host species, and white and grey nodes represent the two parasitoid species. Solid arrows show possible trophic interactions, and dashed arrows show possible competitive interactions in each treatment. The type of competition between host species (intraspecific/interspecific) and presence or absence of parasitoids in the cages were manipulated in a fully factorial design: a) intraspecific competition, b) interspecific competition, c) parasitism only, and d) all interactions

The experimental temperatures were chosen to simulate current mean yearly temperature at the two study sites [47]: 23.2 ± 0.4°C (65.9 ± 2.8% humidity), and projected temperatures representing a plausible future scenario under climate change: 26.7 ± 1.0°C (65.1 ± 2.8% humidity)[52]. The simulated difference was therefore 3.5°C. Vials were placed at their corresponding temperature treatment from the first day the adult hosts were introduced for mating and oviposition to the last emergence (up to 40 days).

To calculate parasitism rates for each host-parasitoid species pair, pupae from the three vials of each box were randomly sampled 12 days after the initiation of the experiment. All sampled pupae were transferred into one or two 96-well PCR plates (on average 169 ± 30 s.d. pupae sampled per box) and kept at their corresponding temperature treatment until adult insects emerged (up to 40 days for the slowest-developing parasitoid species). Sampled pupae were identified to their corresponding host species, and the outcome was recorded as either a host, a parasitoid, an empty pupal case, or an unhatched pupa. We assumed that any pupae which were empty at the time of sampling resulted in adult hosts because this period was too short for parasitoids to complete development and emerge. We calculated parasitism rates from the pupae sampled in plates only. Parasitism rates were calculated as the proportion of each parasitoid species that emerged from the total number of sampled pupae of each host species. Attack rates were not calculated because the exact initial numbers of host larvae available for the parasitoids were unknown.

All hosts that emerged (from both vials and sampling plates) were used to quantify the following aspects of host community structure: abundances of each host species, and their frequencies (i.e., the fraction of all host individuals belonging to each host species). All hosts and parasitoids that emerged from vials before and after subsampling for parasitism rates were collected, identified, and stored in 95% ethanol until the second generation started emerging from the vials (i.e., hosts were no longer collected after four days without any emergence). Effect of treatments on mean host body mass were also investigated, as an increase in temperature generally produces smaller individuals, which could influence the outcome of competition [27]. Individual dry body mass of hosts was measured with 1 μg accuracy using a Sartorius Cubis ™ micro-balance. Only fully-eclosed and intact individuals were included in measurements.

### Statistical analysis

All replicates with fewer than ten total emergences or pupae were removed from analyses of host abundances, frequencies, and parasitism rates (Supplementary Table S2), as these outcomes were associated with low success during the mating process and not with experimental treatments (results with the whole dataset can be found in Supplementary Table S3 online). Data were analyzed with generalized linear models (GLMs). After testing for overdispersion of the residuals, abundance data were modeled using a negative binomial error distribution, host body mass using a gaussian error distribution, and frequencies of host species and parasitism rates using a quasibinomial error distribution. Parasitism (two levels), type of competition (two levels), host species (three levels), parasitoid species (two levels), and temperature (two levels) were included as categorical predictor variables within each model. Blocks were included in the models as a fixed effect because of the small number of blocks. Each two-way interaction was tested and kept in our models if judged to be significant on the basis of backward selection using ANOVA likelihoodratio tests. Interaction between temperature and parasitism, temperature and competition, and parasitism and competition were systematically kept in our models as the experiment was designed to test for the significance of these interactions. The three-way interaction between temperature, parasitism, and competition was tested for host abundances, host frequencies, and host body mass, but was not significant. Significance of the effects was tested using type III analysis of deviance with F-tests. Factor levels were compared using Tukey’s HSD *post hoc* comparisons of all means, and the *emmeans* package. Model assumptions were verified with the *DHARMa* package. All analyses were performed using R 3.5.2[53] with the packages *stats,MASS*[54], *car*[55],*performance[56], DHARMa[57],* and *emmeans[58].*

## Results

In total, 7627 individuals (7063 hosts and 564 parasitoids) were reared across all treatments and replicates (238.3 ± 13.3 s.d. on average per box). Across all treatments and replicates, 2717 pupae were sampled in total for estimating parasitism rate, of which 2227 (82%) produced an adult host or parasitoid. Mean host abundances, host body mass, and parasitism rates are presented for each treatment in Supplementary Table S4 online. We focused on the effects of temperature, parasitism, competition and their interactions on host abundances, host proportions, and host body mass (Table 1).

**Table 1:** ANOVA table showing the effect of temperature (23°C or 27°C), parasitism (yes or no), competition between host species (intraspecific or interspecific), host species (n = 3), parasitoid species (n = 2), interactions between terms, and block (n = 4) on host abundances, host frequencies, host body mass, and parasitism rate. Degrees or freedom (Df) for each F-ratio are given for each factor and for the error. F values are presented with the significance of the effect: (***) P < 0.001, (**) P < 0.01, (*) P < 0.05, (.) P < 0.1, (ns) P > 0.05.

|  | Df | Host abundance |  | Host frequency |  | Host body mass |  | Parasitism rate |  |
| --- | --- | --- | --- | --- | --- | --- | --- | --- | --- |
| **Temperature** | 1 | 1.41 | (ns) | 0.47 | (ns) | 1.88 | (ns) | 4.89 | * |
| **Parasitism** | 1 | 21.80 | *** | 0.03 | (ns) | 2.98 | (.) | - | - |
| **Competition** | 1 | 0.15 | (ns) | 0.06 | (ns) | 10.76 | ** | 1.14 | (ns) |
| **Host species** | 2 | 27.07 | *** | 64.7 | *** | 426.64 | *** | 2.47 | (.) |
| **Parasitoid species** | 1 | - | - | - | - | - | - | 2.29 | (ns) |
| **Temperature x Parasitism** | 1 | 0.26 | (ns) | 0.05 | (ns) | 0.60 | (ns) | - | - |
| **Temperature x Competition** | 1 | 0.00 | (ns) | 0.07 | (ns) | 1.32 | (ns) | 0.04 | (ns) |
| **Temperature x Host species** | 2 | 7.90 | *** | 24.12 | *** | - | - | - | - |
| **Parasitism x Competition** | 1 | 1.58 | (ns) | 0.00 | (ns) | 4.49 | * | - | - |
| **Competition x Host species** | 2 | - | - | - | - | 27.80 | *** | - | - |
| **Host x parasitoid species** | 2 | - | - | - | - | - | - | 20.23 | *** |
| **Block** | 3 | 1.02 | (ns) | 0.47 | (ns) | 4.53 | ** | 1.49 | (ns) |
|  |  |  |  |  |  |  |  | - |  |
| **Df error** |  | 68 |  | 68 |  | 65 |  | 70 |  |
| **R<sup>2</sup>** |  | 0.87 |  | 0.05 |  | 0.93 |  | 0.10 |  |

### Direct effect of warming on the host community

The effect of temperature on host relative abundances varied significantly across host species (Table 1, Figure 2). At 23°C, *D. birchii* and *D. pseudoananassae* had similar relative abundances across treatments (mean frequency of *D. birchii* = 0.426 ± 0.05; mean frequency of *D. pseudoananassae* = 0.471 ± 0.05). At 27°C, host community was dominated by *D. pseudoananassae* for all treatments. *D. pseudoananassae* relative abundances increased by 12.8% while *D. birchii* relative abundances decreased by 56.1% (mean frequency of *D. birchii* = 0.187 ± 0.02; mean frequency of *D. pseudoananassae* = 0.743 ± 0.02). Elevated temperature had no effect on host body mass (F_1,65_ = 1.88, P = 0.175, Supplementary Figure S5).

**Figure 2.**
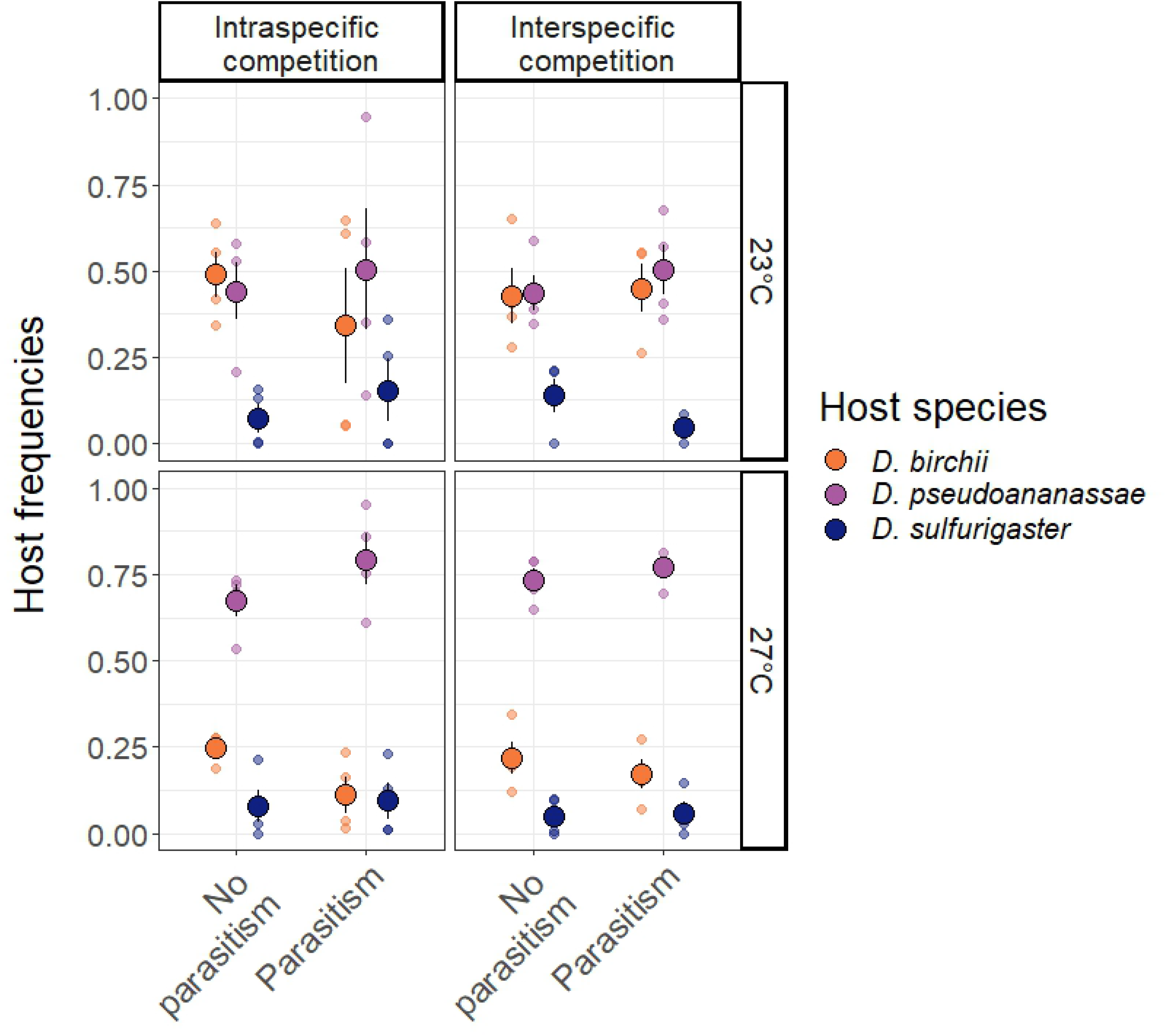
Effect of experimental treatments on host proportions. Experimental warming changed the relative proportions of hosts for all treatments. See Figure 1 for detailed description of the treatments. The small points represent the values from each block, the large points represent the grand mean, and the bars represent standard errors of the means.

### Effect of biotic interactions on the host community

Parasitism significantly reduced mean abundances of all three host species by 50 ± 0.22 (s.e.m.) hosts on average across species (β = −0.339, F_1,68_ = 21.80, P < 0.0001; Figure 3a), and the negative effect of parasitism was consistent across host species (Table 1). Competition type did not significantly impact host abundances or relative host frequencies. Effects of competition on host body mass depended both on host identity (F_2,65_ = 27.80, P < 0.0001), and on presence or absence of parasitoids (F_1,65_ = 4.87, P = 0.038). *D. pseudoananassae* was the host species that varied the most in body mass with treatments (Supplementary Figure S5). Its body mass decreased with interspecific competition in the absence of parasitoids, but increased with interspecific competition with presence of parasitoids. Changes in body mass for the other two host species were less pronounced.

**Figure 3.**
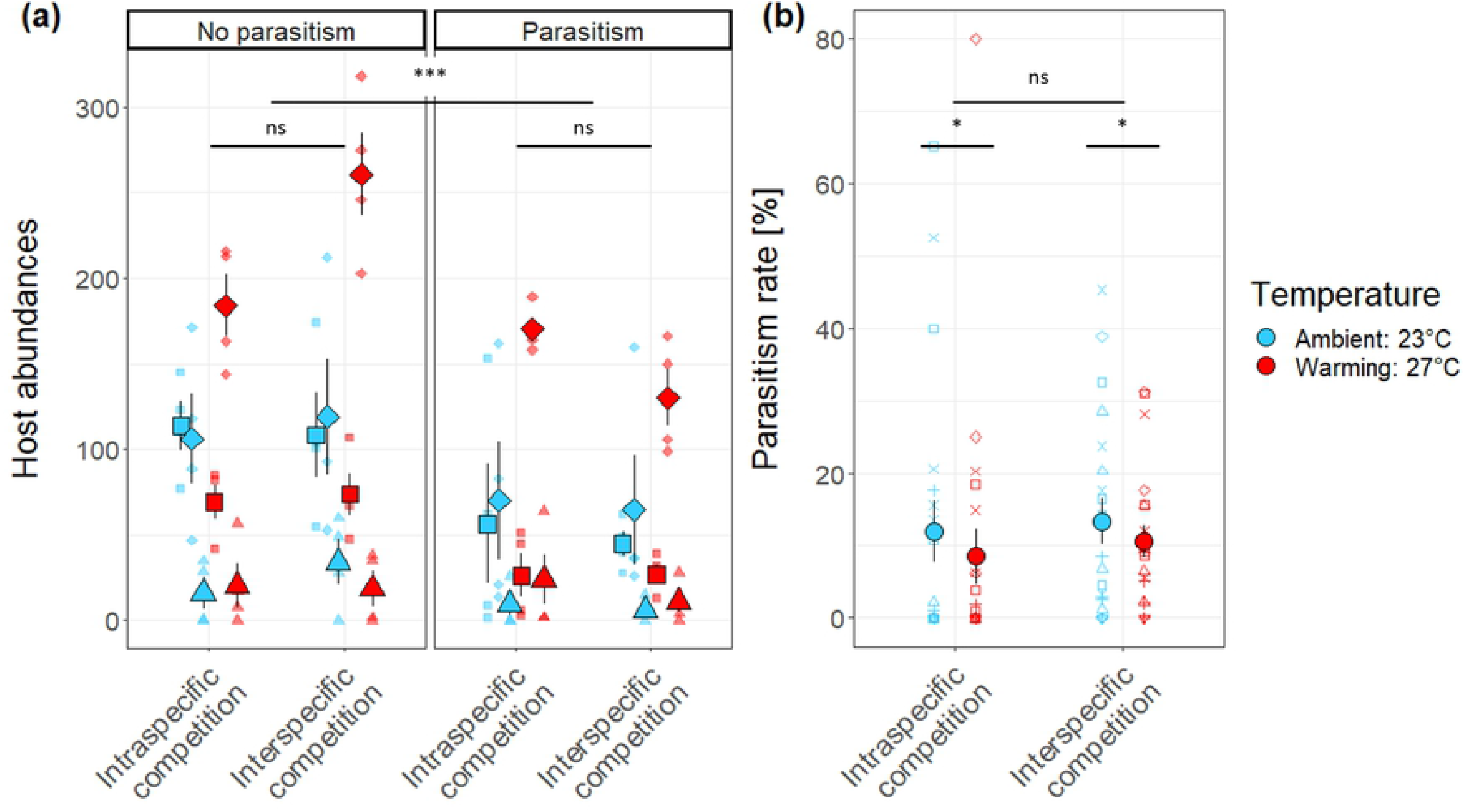
Effect of experimental treatments on host community and host-parasitoid interactions. (a) Host abundances (■: *D. birchii,* ♦: *D. pseudoananassae,* ▲: *D. sulfurigaster)* were significantly reduced by parasitism across treatments. (b) Parasitism was reduced at higher temperature (□: *Asobara* sp. – *D. pseudoananassae*, ^: *Asobara* sp. – *D. birchii,* +: *Asobara* sp. – *D. sulfurigaster,* × *Leptopilina* sp. – *D. pseudoananassae*, ◊: *Leptopilina* sp. – *D. birchii,* V: *Leptopilina* sp. – *D. sulfurigaster).* See Figure 1 for detailed description of the treatments. The small points represent the values from each block, the large points represent the grand mean, and the bars represent standard errors of the means. Significance of treatment effects is indicated as follows: (***) P < 0.001, (**) P < 0.01, (*) P < 0.05, (.) P < 0.1, (ns) P > 0.05

### Indirect effect of warming on host community structure through parasitism and interspecific competition

Experimental warming significantly decreased parasitism for all host-parasitoid pairs (β = −0.29, F1,70 = 4.89, P = 0.030, Table 1, Figure 3b). However, temperature did not interact significantly with either parasitism or competition in affecting any of our measures of community structure (P > 0.05, Table 1), suggesting that the relative effects of parasitism and competition as structuring agents were not modified by experimental warming.

## Discussion

Our experiment revealed that experimental warming directly affected host community structure through differences in thermal performance among species, and decreased parasitism rates. However, warming did not impact the effect of parasitism on host community structure over the timescale investigated.

Our results suggests that ongoing rises in global temperatures could directly alter host community structure through differences in thermal performance across species, as has been shown for communities of fish [59], plants [60], and insects [61]. Changes in host frequencies in warmer temperatures was primarily due to a dramatic increase in the relative abundance of a single host species, *D. pseudoananassae*, the species with the largest thermal performance breath [48]. This increase occurred across all combinations of parasitism and competition treatments, and without a change in *Drosophila* body mass, suggesting a direct effect of temperature on host fecundity due to the preferred temperature of the adults for egg-laying and/or offspring egg-to-adult viability related to their optimal temperature [62]. Surprisingly, the dominant species at warmer temperature, *D. pseudoananassae*, was not the one with the highest thermal performance optimum measured by MacLean *et al.* [48]. However, in our system, its distribution is limited to low elevation sites, and this species has a higher thermal tolerance than either of the other two species considered [48]. In nature, *Drosophila* species distributions are driven by differences in innate thermal tolerance limits, with low phenotypic plasticity for thermal tolerance limits in both widespread and tropical species [49]. This suggests that warming temperatures, in the context of global climate change, will have a strong effect on community composition through direct effect on fitness.

Our data also revealed a significant decrease in parasitism rates with warming. Reviews suggest that parasitism would decrease under global warming scenarios due to an increase in parasitoid mortality, and host-parasitoid spatial and temporal asynchrony [13,43]. However, presence of parasitoids significantly decreased abundances of the three host species independently of the temperature regime, suggesting that warming treatments did not decrease attack rate, but decreased parasitoid virulence. This experiment was performed over a single generation, so long-term consequences of decreased parasitism rates with elevated temperatures for host-parasitoid dynamic cannot be assessed, but a decrease in parasitism rates could lead to the release of hosts from top-down control. However, in the case of a simple linear tritrophic interaction, the results of Flores-Mejia *et al.* [63] suggest that parasitoid top-down control might be less sensitive to temperature than previously thought. Moreover, host immune responses might also be sensitive to temperature [64], increasing or decreasing host vulnerability to parasitoid attacks. Therefore, host immune function response to temperature should be considered alongside host thermal performance and tolerance to predict the effects of warming temperature on host communities [13].

Our results demonstrate that differences in thermal performance across host species may be a stronger determinant of how host communities respond to warming temperatures than shifts in the strength of biotic interactions. We used high, but realistic levels of competition and parasitism that would have allowed us to detect their effects on host species relative abundances if they were any. Aspects of our results contrast with those from a field transplant experiment on two species drawn from the same Australian *Drosophila*-parasitoid community [65]. Investigating fitness of *D. birchii* and *D. bunnanda* along an elevation gradient, the authors found an interacting effect between the abiotic environment and interspecific competition. However, the field experiment excluded parasitoids, and the elevational gradient studied is likely to include variations such as humidity as well as temperature, which might influence the outcome [66].

Our study serves as example of the mechanisms that can be expected to drive community responses to global warming, but general conclusions on the potential impact of warming temperature on host-parasitoid networks will require replication with different species compositions and different systems. Especially, most host-parasitoid systems are tri-trophic (plants-arthropods-parasitoids), and climate warming is likely to impact host-parasitoid networks through bottom-up effects [67]. Few such experiments have been undertaken, despite the need to better disentangle direct and indirect effects of warming temperature on species communities. Ideally, future studies will also need to investigate the longer-term dynamics of such systems. Moreover, as temperatures continue to increase, species from diverse taxa are shifting their distribution worldwide to higher latitudes and elevations [68], changing their biotic environment with novel species interactions and different community assemblages [69]. Dispersal was not permitted in this study, but is likely to mediate some of the effects of warming temperature on species and their interactions [29,70].

Understanding the mechanisms driving community responses to warming scenarios is particularly important for tropical communities, which face more severe impacts of climate warming than temperate communities. Here, we demonstrate that warming had a direct effect on our focal tropical *Drosophila* host community through differences in thermal performance, without affecting the relative strength of parasitism and competition. The role of parasitoids as essential top-down control agents of insect populations was not reduced under experimental warming, but parasitism rate decreased, suggesting that an indirect effect of warming temperature on the structure of host community through the effect of parasitism could be observed after several generations.

## Acknowledgements

We thank Inga Freiberga, Anna Jandová, Martin Libra and Joel Brown for their help during the collection of the data and Petr Šmilauer for his advices on the statistical analysis. The drawings used for Figure 1 was made by Tereza Holicová.

## Supplementary Information captions

**Supplementary Table S1.** Mean number of offspring per species with 10, 30, 60, 90 or 180 adult hosts (1:1 sex ratio) in 5 mL host-media glass vial. Choice of host number in the main experiment design was based on these preliminary data to correspond to strong competition for all host species

**Supplementary Table S2.** Number of observations per temperature, treatments (Intraspecific competition: no interaction between host species, Interspecific competition: direct competition between host species, Parasitism: parasitoids present without direct interaction between host species, All interactions: both presence of parasitoids and interspecific competition between host species), and host species in the whole dataset, and with the reduced dataset used for analyses (excluding observations with fewer than 10 emerging insects or pupae)

**Supplementary Table S3.** ANOVA table for the whole dataset showing the effect of temperature (23°C or 27°C), parasitism (yes or no), competition between host species (intraspecific or interspecific), host species (n = 3), interactions between terms, and block (n = 4), on host abundances, and host frequencies. Degrees or freedom (Df) for each F-ratio are given for each factor and for the error. F values are presented with the significance of the effect: (***) P < 0.001, (**) P < 0.01, (*) P < 0.05, (.) P < 0.1, (ns) P > 0.05

**Supplementary Table S4.** Summary table for mean (± s.d.) host abundances (Host ab.), individual host body mass (Host BM), total parasitism rate (PR), and parasitism rates of each parasitoid species *(Asobara* sp. and *Leptopilina* sp.) for each temperature (23 and 27°C), treatments (competition: intra or inter, parasitism: yes or no), and host species *(D. birchii, D. pseudoananassae, D. sulfurigaster)*

**Supplementary Figure S5.** Interactive effect of competition with host species, and with presence of parasitoids on mean host body mass (squares: *D. birchii*, diamonds: *D. pseudoananassae*, triangles: *D. sulfurigaster).* See Figure 1 for detailed description of the treatments. The small points represent the values from each block and each host-parasitoid pair, the large points represent the grand mean, and the bars represent standard errors of the means. Blue: ambient temperature (23°C), red: warming treatment (27°C)

